# Ionizing Radiation Shapes Genome Evolution Through Nuclear Abnormalities That Trigger Delayed Proliferative Death

**DOI:** 10.64898/2025.12.01.691539

**Authors:** Mian Zhao, Bejo Presila, Cheng-Zhong Zhang, Spektor Alexander

**Author notes:** Co-first authors (Mian Zhao, Bejo Presila).

## Abstract

Ionizing radiation (IR) is widely used in cancer therapy, yet the mechanisms by which IR-induced DNA damage translates into delayed proliferative death and secondary malignancy risk remain incompletely defined. Radiation generates double-strand breaks (DSBs) that are frequently misrepaired, giving rise to micronuclei and chromosome bridges—abnormal nuclear structures known to drive chromosomal instability. Here, a modified live-cell imaging and single-cell whole-genome sequencing (WGS) approach was used to directly connect early cytological responses to genome-wide outcomes in cells exposed to graded IR doses. Tracking asynchronously cycling cells revealed that micronuclei, chromosome bridges, and nuclear fragmentation form at high frequency during the first mitosis after irradiation, in a strongly cell-cycle–dependent manner: micronuclei were most frequent after S-phase irradiation, whereas chromosome bridges predominated when damage occurred in G1 or S phase. Lineage tracing combined with clonogenic assays showed that formation of abnormal nuclear structures in the first post-irradiation division is a major determinant of long-term proliferative capacity, with centromere-containing (centric) fragments and bridges exerting a stronger anti-proliferative effect than acentric micronuclei. Single-cell WGS of daughter and granddaughter cells demonstrated that these structures generate reciprocal copy-number changes, clustered rearrangements, and chromothripsis-like events that are largely incompatible with sustained proliferation. In contrast, bulk WGS of surviving single-cell–derived clones revealed only modest increases in single-nucleotide variants and indels but an enrichment of nonhomologous end-joining–associated deletion signatures and selected structural variants, including amplification of a pre-existing 12p gain. These findings identify abnormal nuclear structures as key intermediates linking IR-induced DNA lesions to mitotic catastrophe and provide insight into selective tolerance of specific rearrangements.

## Introduction

Radiation therapy (RT) is among the most widely used modalities in cancer treatment, employing ionizing radiation (IR) to eradicate malignant cells while minimizing collateral damage to surrounding normal tissue^1,2,3^. When IR penetrates biological tissue, it induces radiolysis of water molecules in the cytoplasm, generating reactive oxygen species (ROS) that chemically damage cellular macromolecules. Among these, DNA is the most critical target, with ROS-mediated lesions leading to single- and double-strand breaks (DSBs) that trigger complex repair responses^4,5^.

Despite its clinical efficacy, RT carries long-term risks. Years after treatment, some patients develop secondary malignancies—most often high-grade sarcomas with poor clinical outcomes^6^. These neoplasms typically originate from normal tissues exposed to therapeutic radiation^7,8^, where cumulative IR-induced mutations are thought to drive oncogenic transformation.

Although secondary malignancies pose substantial clinical challenges, the genomic evolution of irradiated cells remains incompletely defined. Existing studies indicate that IR exposure generates a dose-dependent increase in DSBs, which are frequently repaired via the error-prone nonhomologous end-joining (NHEJ) pathway^9,10^. This process produces characteristic mutational signatures, such as microhomology-mediated deletions and balanced inversions^11,12^. A subset of irradiated cells may bypass cell-cycle checkpoints despite harboring unrepaired DNA damage, propagating DNA damage and promoting chromosomal instability^13,14^. Consequently, the efficiency and fidelity of the DNA damage response (DDR) and repair pathways are pivotal in determining the ultimate fate of irradiated cells^15^.

The mechanisms translating DNA damage into clonogenic cell death are diverse and include apoptosis, necrosis, necroptosis, ferroptosis, and autophagy-dependent pathways^16,17^. In solid tumors such as carcinomas and sarcomas, however, RT more commonly induces mitotic catastrophe—a delayed, proliferation-associated form of cell death wherein progeny cells progressively lose reproductive capacity over several divisions^18,19,20,21^.

The mechanism by which mitotic catastrophe occurs is incompletely understood, but one possible driver of mitotic catastrophe appears to be the formation of IR-induced nuclear abnormalities, including micronuclei (MN) and chromatin bridges. These aberrations are widely recognized cytogenetic markers of radiation exposure and genomic instability^22,23^. MN arise when acentric chromosome fragments or lagging chromosomes fail to incorporate into daughter nuclei during mitosis, while chromatin bridges typically originate from dicentric chromosomes produced by misrepaired DSBs and inappropriate recombination^24,25^. Both structures can sustain extensive DNA fragmentation, leading to chromothripsis-like complex genome rearrangements, copy-number alterations, and kataegis on affected chromosomes^26,27,28,29,30^.

To dissect how these nuclear abnormalities influence the genomic evolution and proliferative potential of irradiated cells, we developed an approach that integrates live-cell imaging with single-cell genome sequencing and whole-genome sequencing of single-cell–derived clones exposed to different radiation doses. This strategy enables us to determine the cell-cycle stage at the time of irradiation, temporally track individual cells through successive divisions, identifying MN and chromosome bridge formation, while connecting these phenotypes to downstream genome-wide alterations. By linking early cytological responses to genetic outcomes, this strategy provides a direct view of how IR-induced DNA damage and repair shape the long-term genomic trajectories of surviving cells.

## Results

### Radiation induces abnormal nuclear structures

To understand how early cytological responses following irradiation affect genome evolution and proliferation capacity, we employed a modified Look-Seq approach^26^ that combines live cell imaging and whole genome sequencing that enables us to determine individual cell’s cell cycle stage at the time of irradiation in asynchronous population without pharmacologic synchronization thereby avoiding drug-induced confounding stress^29,30^, and then temporally follow the cells through the first several cell cycles to detect formation of abnormal nuclear structures such as MN and chromosome bridges. Individual cell progeny could then be isolated for single-cell whole genome sequencing. Asynchronous RPE-1-hTERT cells with CRISPR/Cas9 knockout of p53, stably co-expressing two fluorescent markers -- GFP-BAF, which allows detection of nuclear abnormalities such as MN, chromosome bridges and primary nuclear envelope ruptures, and mCherry-PCNA, which facilitates identification of the cell cycle phase, were imaged starting one hour prior to irradiation for 72 hours. (Figure 1a). Cell cycle phase at the time of irradiation was determined by mCherry-PCNA status on the image captured just prior and immediately following irradiation. Presence of PCNA foci indicated S phase; if foci were absent (cells in either G1 or G2 phase), cell cycle phase was determined by whether the cell then proceeded into S phase (for G1) or mitosis (for G2). Nuclear abnormalities (MN, chromosome bridges, primary nuclear ruptures, and nuclear fragmentation (defined as ≥3 distinct nuclei)) were detected over several cell divisions during GFP-BAF. To link cytological phenotypes to genomic consequences, daughter or granddaughter cells were individually isolated and subjected to single-cell whole genome sequencing (Supplementary Fig. 1a).

**Fig. 1.**
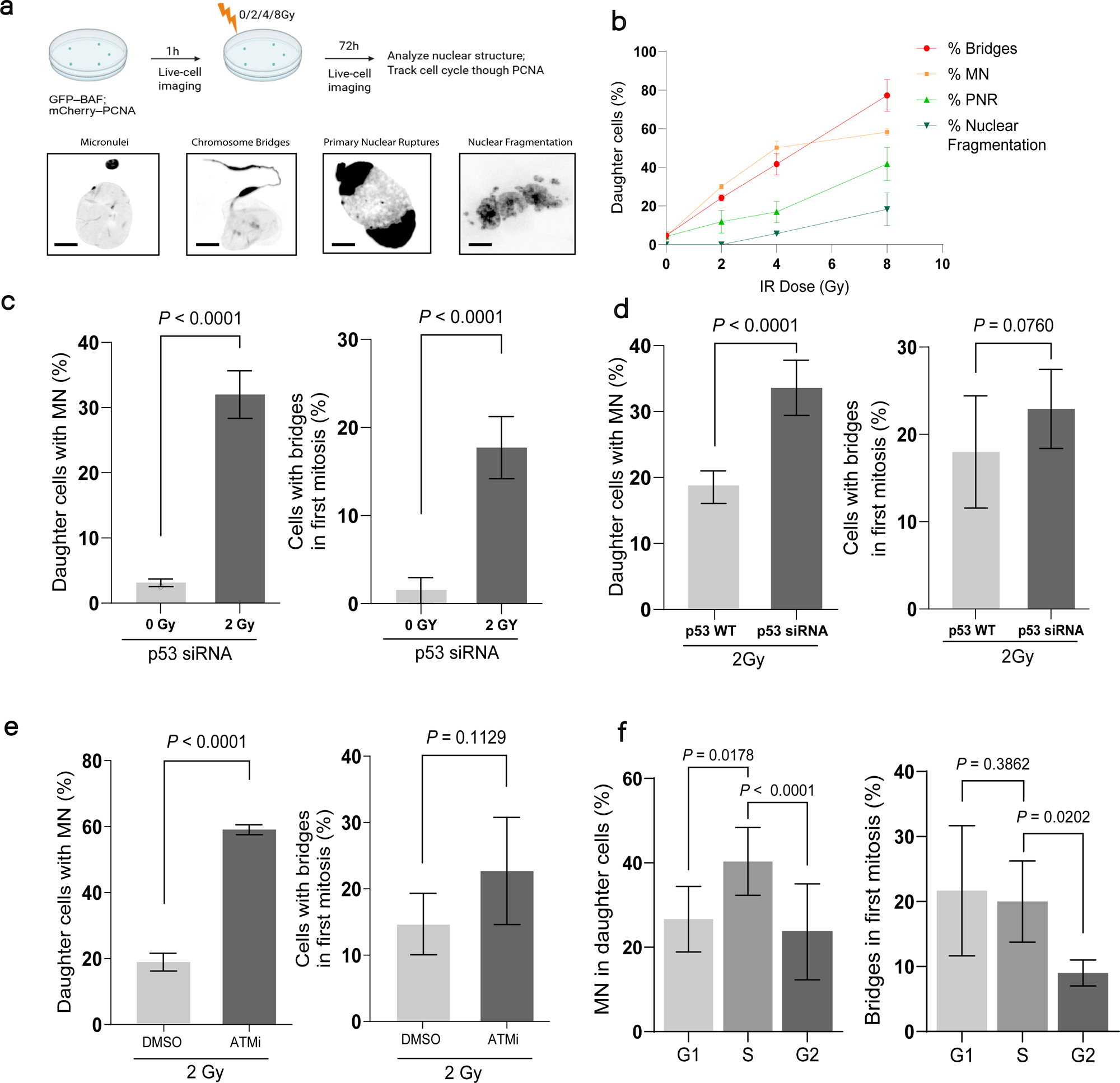
Radiation induces abnormal nuclear structures in the first generation. **(a)** Schematic depicting experimental outline. RPE-1 p53-null cells expressing GFP-BAF and PCNA-mCherry were plated 24 hours before irradiation. Time-lapse confocal imaging began 1 hour pre-IR and continued for 72 h after exposure to 0, 2, 4, or 8 Gy, followed by downstream image analysis. *Bottom*: Representative images of the mitotic errors identified by GFP-BAF. Scale bar, 5 µm. **(b)** Quantification of the percentage of first-generation cells exhibiting the indicated nuclear abnormalities at 0, 2, 4, and 8 Gy. MN - micronuclei, PNR-primary nuclear rupture. Graphs represent mean ± SEM; n = 2 independent experiments. **(c)** Percentage of first-generation daughters with MN (*left*) and chromosome bridges (*right*) in p53-null cells, 0 Gy vs 2 Gy. (**d**) Percentage of first-generation daughters with MN (left) and chromosome bridges (right) at 2 Gy, p53 WT vs p53-null. (**e**) Percentage of first-generation daughters with MN (*left*) and chromosome bridges (*right*) in p53-null cells at 2 Gy, DMSO vs ATMi. (**f**) Percentage of first-generation daughters with MN (*left*) and chromatin bridges (*right*) in RPE-1 p53-null cells, irradiated with 2Gy at cell cycle phases as indicated. For (**c–f**), mean ± SEM (n=3 independent experiments). P-values by unpaired two-tailed Student’s t-test.

As previously reported^31^, we observed a high number of nuclear abnormalities following progression through mitosis even at low radiation doses (2Gy), with roughly 25% increase in micronucleation and 20% increase in chromosome bridge formation from 0 to 2Gy (Figure 1b). Micronucleation, chromosome bridge formation and primary nuclear ruptures increased with increasing radiation dose in a roughly a linear fashion, with 75% of daughter cells exhibiting chromosome bridges by 8Gy. Above 4Gy, we observed an increasing number of cells with nuclear fragmentation (≥3 distinct nuclei or micronuclei), with corresponding levelling off of cells exhibiting 2 or fewer MN.

We next proceeded to identify the relationship between checkpoint signaling and formation of abnormal nuclear structures. We determined the interval from irradiation to first mitosis at different doses, and found that the interval increased from 9 h at 0 Gy to 27 h at 8 Gy consistent with activation of DNA damage checkpoint (Supplementary Fig. 2b). At higher doses, increasing numbers of cells did not enter mitosis by the end of the video, suggesting possible senescence. We further assessed the effect of p53 status and ATM inhibition on MN. p53-null and ATM-inhibited (ATMi) cells exhibited significantly higher rates of micronucleation and chromatin bridge formation relative to controls upon progression through the first mitosis following irradiation with 2Gy (Fig. 1d,e), consistent with the key roles of p53 and ATM in cell cycle checkpoint and DNA response initiation respectively. In the absence of these factors, higher numbers of cells get to first mitosis with unrepaired DNA damage, leading to formation of nuclear abnormalities including MN and chromosome bridges.

Previous studies showed that micronuclei and chromosome bridges can drive complex genome rearrangements and chromothripsis^25,26,27,29,30^. To test whether IR-induced abnormal nuclear structures have similar consequences, we isolated daughter cell pairs performed following progression through mitosis and performed WGS on individual daughter cells. We observed reciprocal copy number (CN) alterations (segmental gains in the daughter that inherited MN or chromosome bridge segments with reciprocal CN loss in the sister cell) in the first cell cycle following formation of these structures (Supplementary Fig. 1b). We also isolated individual granddaughter cells following progression of MN or bridged cells through the subsequent mitosis and observed clustered complex chromosomal rearrangements and chromothripsis that correspond to the regions of segmental gain^25,26,27,29,30^ (Supplementary Fig. 1c). These results indicate that radiation-generated micronuclei and chromosome bridges result in genomic consequences comparable to abnormal nuclear structures induced by other methods such as nocodazole and Mps1 inhibitor, which result in whole chromosome mis-segregation through formation of merotelic attachments and CRISPR/Cas9 targeted cuts, which produce acentric MN and chromosome bridges through sister chromatid fusion.

### Formation of abnormal nuclear structures is cell cycle-dependent

Our experimental system that does not depend on drug synchronization allows us to determine whether the frequency of MN and chromosome bridge formation depends on the cell cycle phase at the time of irradiation. We observed a significantly higher rate of chromosome bridge formation when the cells were irradiated in G1 or S phase of the cell cycle, and lower frequency in G2 (Figure 1f). Because photon radiation causes DSBs predominantly on a single chromatid, DSBs produced in G1 phase or in S phase prior to replication of a given region would lead to broken ends on both sister chromatids upon completion of DNA replication, this resulting in a high rate of sister chromatid fusion, In contrast, irradiation in G2 results in only a single broken chromatid, with the low rate of sister chromatid fusion and chromosome bridge formation. We also observed a higher rate of MN formation when the cells were irradiated in S phase compared to G1 and G2 (Figure 1f). We postulate that this may occur due to the interaction between single-strand breaks and base damage, the more common forms of DNA damage induced by IR than DSBs^34,35^, and DNA replication forks, which result in formation of one-ended DSBs and thus higher probability of unrepaired DSBs upon mitotic entry. Taken together, these results support that both MN and chromosome bridge formation are cell cycle-dependent, with highest probability of forming a MN when a cell is irradiated in S phase and highest frequency of chromosome bridges in G1 and S.

### Abnormal nuclear structures are a major determinant of cell’s proliferation capacity

Radiation often does not kill cells immediately; instead, it triggers delayed cell death^18,19,20,21^, but the precise mechanism by which this occurs remains unclear. Because formation of even a single MN or chromosome bridge can dramatically restructure the genome, we hypothesized that formation of MN or chromosome bridges are the pain source of proliferative cell death following irradiation. To test this hypothesis, we assessed whether formation of MN or chromosome bridges in the first generation following irradiation affects ability of cells to grow into clones. We sorted a single cell per well into 384-well plates, irradiated at 2 Gy, and followed individual cells by live cell imaging for 24–48 hours to detect MN and bridge formation upon mitotic progression (Figure 2a). We then cultured the plates for 8 days and counted the number of cells present in each well at the end of incubation period. Most of the cells that did not develop MN or bridges in the first generation following mitosis were able to form clones of approximately 200 cells on average. In contrast, the majority of the cells that formed MN or chromosome bridges stopped proliferating after several cell divisions and failed to form clones with only two clones reaching 200 cells (Figure 2b). These results demonstrate that formation of abnormal nuclear structures in the first mitosis following irradiation is associated with the loss of proliferative capacity.

**Fig. 2.**
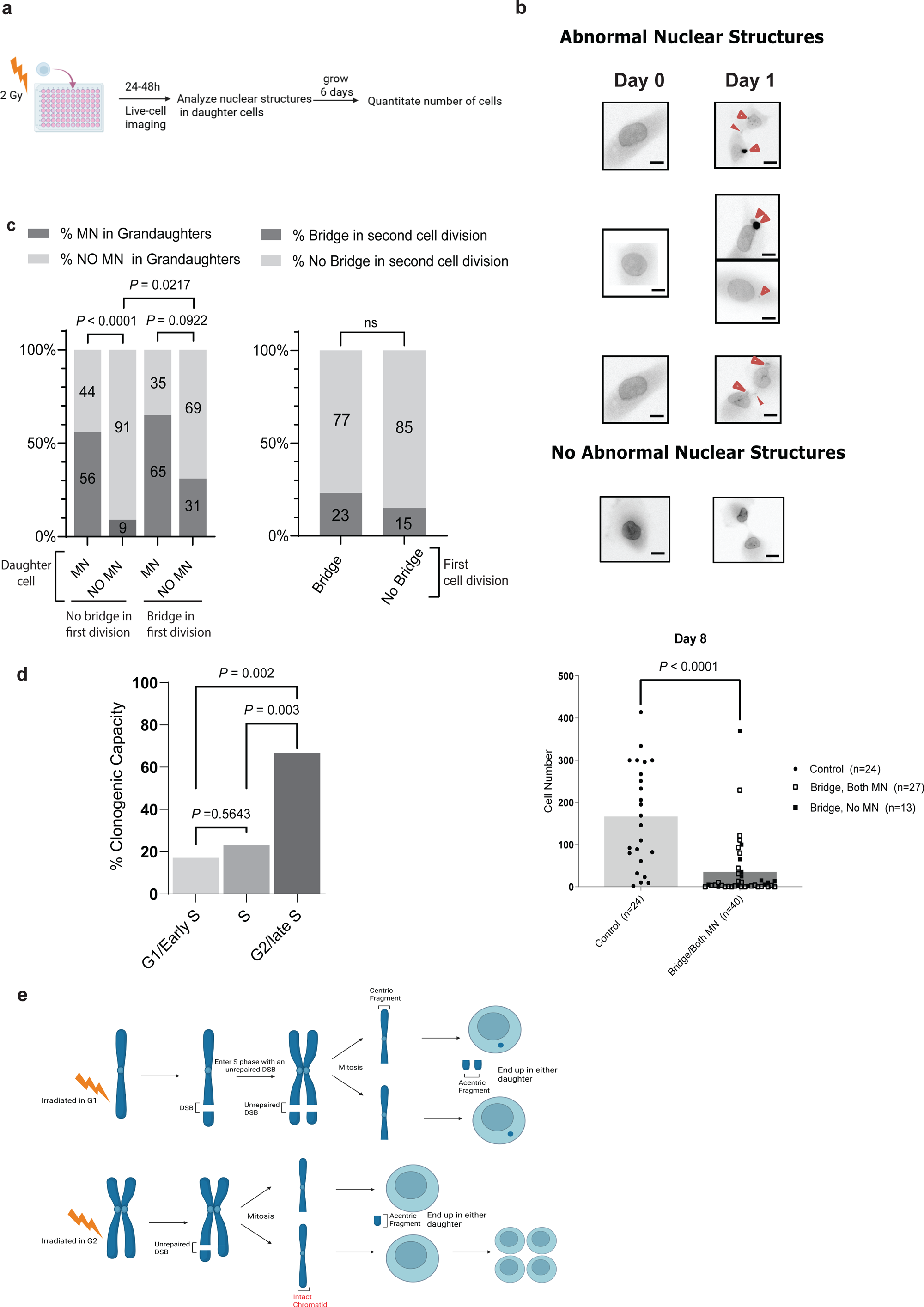
Abnormal nuclear structures are a major determinant of cell’s proliferation capacity. **(a)** Schematic depicting experimental outline. Single cells were plated one per well in 384-well plates, irradiated with 2 Gy, and imaged by live cell imaging for 24–48 hours to identify abnormal nuclear structures that form in the first mitosis. Cells were then returned to the incubator for 6 days, and the total number of cells was determined on day 8. (**b**) *Top:* Representative live-cell snapshots from the experiment in **a** at day 0 and day 1. Scale bar, 5 µm. *Bottom:* Chart depicting number of cells per clone grouped by absence vs. presence of abnormal nuclear structures following first mitosis (*n* = 24 and *n* = 40, respectively). Circle = cells without abnormal nuclear structures; open square = chromosome bridge and both daughter cells with micronuclei; filled square = chromosome bridge without micronuclei. Bars indicate mean. (**c**) *Left*: Percentage of granddaughter (second generation) cells with MN, stratified by whether the first-generation daughter had MN and whether a chromosome bridge formed during the first mitosis. *Right:* Percentage of cells with chromosome bridges during the second mitosis, stratified by whether a bridge occurred in the first division. (**d**) Schematic depicting difference in inheritance of centric and acentric fragments by daughter cells based on the cell cycle phase at the time of irradiation. When cells are irradiated in G1, both sister chromatids exhibit unrepaired DSB following DNA replication, with both daughter cells inheriting the broken centric fragment; in contrast, when a cell is irradiated in G2, only one sister chromatid is broken, and one of the two daughter cells inherits an intact chromatid. (**e**) Percentage of micronucleated or bridged cells that formed a clone after 2 Gy IR at cell cycle phases (G1/S/G2) as indicated. For (**b-d**), mean ± SEM (n=3 independent experiments). P-values by unpaired two-tailed Student’s t-test.

To understand how abnormal nuclear structures limit proliferation, we examined whether abnormal structures are transmitted across multiple cell cycles. Using live cell imaging, we tracked single cells across two mitoses and quantified the intergenerational transmission of MNs and chromosome bridges. We observed that 56% of granddaughters from MN-positive daughters retained at least 1 MN, versus 9% when daughters were MN-negative (Figure 2c, *left*); the remaining 44% of cell lineages lost MNs after a second mitosis, likely due to reincorporation into the primary nucleus, fragmentation into smaller fragments that are no longer detectable by BAF or degradation. Presence of chromosome bridges in the first generation following irradiation led to increased MN formation in the subsequent generation consistent with chromosome bridge breakage^25,27^. Presence of chromosome bridges in the first generation also increased the incidence of bridges in the next generation, albeit modestly (Figure 2c, *right*), suggesting that breakage-fusion-bridge (BFB) cycles do not necessarily occur in every generation.

Together, these results demonstrate that abnormal nuclear structures persist across multiple cell divisions leading to a cascade of genome instability that ultimately limits proliferation capacity.

### Broken centric fragments limit proliferation capacity

Although most MN/bridge-positive cells failed to grow into clones, a few did form clones after irradiation. The question remains whether this limited proliferation are directly caused by MN or chromosome bridges, and if so, which one? Acentric MN arise due to the mis-segregation of acentric terminal fragments. In contrast, chromosome bridges are formed as a result of centromere-containing (centric) fragments undergoing sister chromatid fusion, forming a dicentric chromosome. To determine whether one of these structures is responsible for limited proliferation, we compared clonogenic capacity in cells that formed MN and chromosome bridges after irradiation in G1 and G2 phases of the cell cycle respectively. As shown in Figure 2d, irradiation in G1 results in two broken sister chromatids following DNA replication in S phase, resulting in sister chromatid fusion in some cases, leading to both daughter cells inheriting a broken centric fragment. In contrast, when cells are irradiated in G2, only one of the two sister chromatids is broken, with one of the daughter cells inheriting an intact chromatid. In contrast, acentric fragments mis-segregate into MN with equal probability whether the cells are irradiated in G1 or G2. If centric fragment is responsible for limiting proliferation of the cells, we expect that G2 irradiated cells that form abnormal nuclear structures will be more likely to grow into a clone since one of the two daughter cells will inherit an intact sister chromatid. Indeed, we observed that over 60% of G2-irradiated cells that formed MN or chromosome bridge following the first mitosis were able to grow into clones in contrast to approximately 20% in G1- or S-irradiated cells. These results suggest that the centromere-containing chromosome fragment, which is prone to undergo breakage-fusion-bridge cycles^36^ and recombination with other chromosomes resulting a cascade of genome instability, is the source of limited proliferation. In contrast, acenric MN are a marker of limited proliferation, but not necessarily the cause of it.

### WGS reveals increased mutational burden in radiation-surviving clones

To understand genomic evolution of cells that proliferate after irradiation, we cultured single-cell derived clones after 0, 2, or 12 Gy of IR and quantified clonogenic survival after irradiation without pre-screening for nuclear abnormalities: approximately 55% of cells formed colonies at 2 Gy, whereas fewer than 1% did so at 12 Gy (Supplementary Fig. 2b). We next isolated single-cell derived cones for each of these dose levels and subjected them to high-depth WGS. We quantified single-nucleotide variants (SNVs) and indels per clone. Since some of the clones were inferred to be tetraploid, we restricted our analyses to diploid clones (n = 29; see methods). Clones derived from high-dose irradiated cells showed a modest increase in *de novo* SNVs and a significant increase in indels relative to low dose unirradiated clones. High dose clones carried about 1,250 SNVs and approximately 200 indels. We calculated mutation frequency on disomic chromosomes, and observed a similar pattern as with *de novo* variant counts. (Figure 3a). Because *de novo* variant counts include mutations acquired during clonal expansion in addition to those caused by radiation, we assessed relative burdens of mutational signatures (Figure 3b). Most SNV signatures did not differ significantly among the three groups, with the highest burden coming from a clock-like signature associated with cell proliferation^37^. By contrast, an NHEJ-associated indel signature (ID8)^37^, characterized by >5-bp deletions and small deletions with microhomology (>1 bp), was enriched in 12 Gy clones (Figure 3b), consistent with prior studies^38^ and the observation that DSBs are repaired to a large extent by classical NHEJ^39^.

**Fig. 3.**
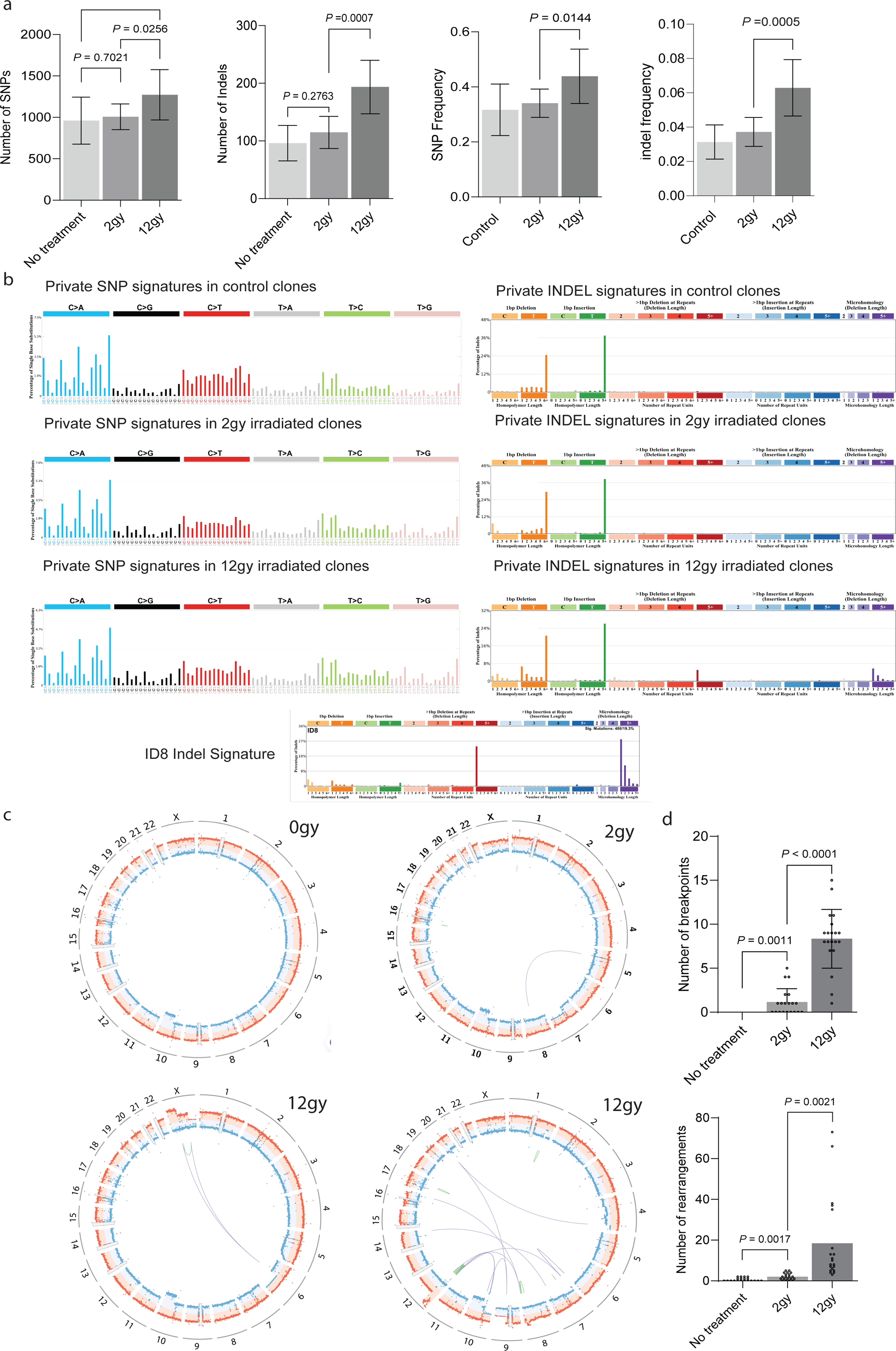
Radiation increases the burden of indels, rearrangement junctions, and genomic breakpoints. (**a**) Quantification of SNVs and indels, with SNV/indel frequencies normalized by chromosome length (Mb) on disomic chromosomes (0, 2, 12 Gy; n = 9, 5, 14). (**b**) Mutational signature spectra of SNVs (*left*) and indels (*right)* generated with SigProfilerExtractor for single-cell derived clones at 0, 2, and 12 Gy (n=9, 5, 14). An indel signature is apparent at higher dose, whereas no distinct SNV signature is detected. **(c)** CIRCOS plots of DNA copy number (90-kb bins) for representative control, 2 Gy, and 12 Gy clones. The blue and orange dots indicate haplotype-specific copy number for the two homologs. Green links mark intrachromosomal rearrangements; orange links mark interchromosomal rearrangements. *Bottom left:* CIRCOS plot showing a complex copy-number state and clustered rearrangements on chr12p from a representative 12 Gy–irradiated clone. (**d**) Quantification of rearrangement junctions and chromosomal breakpoints (0, 2, 12 Gy; n = 20 each). For (**a and d**), mean ± SEM, P-values by unpaired two-tailed Student’s t-test

We next performed breakpoint and structural variant analysis. CIRCOS plots of representative clones for each of the dose groups are shown in Figure 3c. We observed a significant increase in breakpoints in structural variants (SVs) in cells irradiated at high dose (12Gy) compared to low dose (2Gy) and unirradiated cells (Figure 3d). Haplotype-specific copy-number analysis showed a mean of 8 breakpoints per 12-Gy sample (segmental, non-ancestral copy-number changes), versus 1 and 0 at 2Gy and 0 Gy, respectively. SV analysis revealed a significantly increase in rearrangement junctions in single-cell–derived clones irradiated at 12 Gy compared to the other two cohorts. Nevertheless, the total number and complexity of chromosome rearrangements observed even in high dose samples was very low compared to single-cell granddaughter samples irradiated at a much lower dose (2Gy). The majority of high-dose samples had 20 or fewer chromosome rearrangements, with 4 samples having a much larger number of chromosome rearrangements (30-80 rearrangements/cell). Interestingly, these high-SV outliers shared a segmental 12p gain, which is a known subclonal feature of RPE-1 cells, but was further amplified in irradiated cells, suggesting positive selection. Most of the observed rearrangements in these highly rearranged samples occurred in this gained region. Taken together, these results suggest that most chromosomal rearrangements that arise following irradiation are deleterious for cell proliferation and thus are not present in single-cell derived clones. Nevertheless, not all rearrangements are universally deleterious and cells may be able to tolerate rearrangements that confer proliferation advantage due to the presence of oncogenes or presence of extra copies due to pre-existing aneuploidy.

## Discussion

By linking the phenotype observed under the microscope with its direct genomic consequences, our study defines the mechanism by which ionizing radiation limits proliferative capacity of cells and results in a delayed replicative death frequently referred to as mitotic catastrophe. We show that abnormal nuclear structures that form at high frequency during the first mitosis after exposure to IR are the major determinant of cell’s clonal capacity. Because micronuclei and chromosome bridges form at high frequency even after a relatively low dose of radiation (2Gy), they lead to extensive genome rearrangement and a cascade of genome instability that becomes incompatible with further cell proliferation within a few cell cycles as evident from our observation that most complex genome rearrangements seen in the second generation are absent from clones at both low and high doses of IR. In contrast, there is a relatively low burden of SNVs and indels directly caused by radiation even at higher doses of IR consistent with prior studies^38^.

We demonstrate that the formation of these structures is cell-cycle dependent, with the highest burden of MN in S phase likely due to the interaction between active replication forks and single-strand breaks and base damage caused by radiation, leading to fork stalling and higher burden of single-ended DSBs^40^.

Further studies are needed to understand the interaction between different types of radiation-induce damage and its genomic consequences. In contrast, chromosome bridges form less frequently in G2, when DSB affects only a single chromatid and does not result in a high rate of sister chromatid fusions.

While both MN and chromosome bridges can dramatically rearrange genomes, they do not have equal effect on cell proliferation capacity. This is evident from the fact that G2-irradiated cells that form MN and/or bridges are much more likely to form clones than their G1- and early S-phase-irradiated counterparts. We postulate that this is due to the fact that G2-irradiated cells form relatively fewer chromosome bridges and one of the two daughters inherits an intact chromatid. In contrast, MN are equally distributed between the two daughter cells both in G1 and G2. We thus propose that the major driver of replicative death is the centric fragment that remains in the main nucleus and has a high probability to undergo BFB cycles or recombine with other chromosomes, leading to a cascade on genome instability. In contrast, acentric micronuclei induced by radiation are deficient in replication, and re-incorporate into the primary nucleus only at low frequency due to the lack of centromere, thus leading to a high probability of loss over subsequent generations. MN and thus likely a marker of limited replicative capacity, but not the cause of replicative death.

While most genome rearrangements are deleterious for cell proliferation and thus selected against in radiation clones relative to early generation single cells, we observed 4 clones from the high-dose (12Gy) cohort that were able to tolerate a relatively large number of genome rearrangements. Interestingly, in all cases, these rearrangements were predominantly clustered on chromosome 12, which represents an ancestral subclonal gain in RPE-1 cells that was positively selected and amplified in the clones. Two possible explanations exist for the tolerance of these rearrangements. First, this region of chromosome 12 contains RAS oncogene, which promotes proliferation and thus confers a selective advantage. It is also possible that cells in general are able to tolerate rearrangements when extra copies of a chromosome are present. This possibility has important implications, as it suggests that pre-existing aneuploidy, which is present in approximately 90% of all cancers, confers radiation resistance by allowing rearranged cells to proliferate.

In summary, by linking cytological findings with their direct genomic consequences, our study provides important insight into the mechanisms of genome evolution following IR exposure and uncovers nuclear abnormalities as the key determinant of cell’s fate. Detailed understanding of genome evolution following IR is essential not only to optimize therapeutic potential of radiation, but also to understand and prevent the formation of radiation-induced secondary cancers. Future studies will build upon these findings to dissect the specific rearrangement patterns that are tolerated by proliferating cells, and link genomic evolution following IR exposure to the genomic features of radiation-associated secondary malignancies^41,42,43^.

## Methods

### Cell Culture

RPE-hTERT (ATCC) and RPE-1-hTERT p53-null cells (generated by CRISPR–Cas9 knockout, kind gift of David Pellman) were cultured in DMEM/F-12 (1:1; Gibco/Thermo Fisher) supplemented with 10% FBS (Gibco), 100 U/mL penicillin + 100 µg/mL streptomycin (Gibco) at 37 °C in a humidified 5% CO□ incubator.

### ATM inhibition

RPE-1 WT and p53-null cells were pre-treated with the ATM inhibitor (ATMi KU-55933 (Selleck) in DMSO) at a concentration of 10 µM for ∼1 h prior to irradiation. 15 hours following irradiation, cells were washed 5× with prewarmed PBS (37 °C) to wash out the drug, then returned to culture medium.

### Live-cell Imaging and Analysis

1× 10^5 RPE1-hTERT cells stably expressing GFP-BAF and mCherry-PCNA were seeded on a 35-mm □-Dish (Ibidi) 24 h before irradiation. Each dish corresponded to a single radiation dose group (0, 2, 4, or 8 Gy). Time-lapse images were acquired on Nikon Eclipse Ti2 microscope equipped with CSU-W1 spinning disk confocal system, Andor Zyla 4.2P sCMOS camera, NIS-Elements software (Nikon) and OKO Lab incubation chamber (37°C, 5% CO2). Images were acquired every 15 min for 72 h with 7 z-planes at 0.5 μm spacing. Cells were imaged for 1 h prior to irradiation, irradiated and then immediately returned to the microscope to continue live cell imaging. Presence or absence of PCNA foci just prior to and after irradiation was used to determine cell cycle phase at the time of irradiation. Presence of foci prior to and immediately after irradiation indicates the cell was in S phase. If PCNA foci were absent, we determined whether the cell was in G1 or G2 phase by whether the cell enters S phase (by PCNA foci) or mitosis respectively to record PCNA foci. Formation of anormal nuclear structure was assessed using GFP-BAF following progression through mitosis.

### Fixed-cell image and analysis

Cells were fixed in 4% paraformaldehyde (Electron Microscopy Sciences, 15710) for 15 min at room temperature (RT), rinsed 3× in PBS, permeabilized in PBS containing 0.5% Triton X-100 for 5–10 min at RT, washed 3× in PBS, and stained with Hoechst for 20 min at RT. After a final PBS wash, coverslips were mounted in ProLong Gold Antifade (Life Technologies) on glass slides. Imaging was performed on a Nikon Ti2 microscope equipped with a Yokogawa CSU-W1 spinning-disk head using a 60× CFI Plan Apo oil immersion objective (N.A. 1.40, Nikon); z-stacks were acquired at 0.5 µm spacing. Quantitative analysis of nuclear morphology was performed in Image J/Fiji as described in^44^.

### Irradiation Procedures and Generation of Single-Cell–Derived Clones

Cells were plated in 15-cm dishes one day before irradiation at 200 (control), 1,000 (2 Gy), and 10,000 (12 Gy) cells per dish. On the day of counting, cells were washed with prewarmed PBS, trypsinized in 0.05% trypsin-EDTA (Gibco, 25200114), resuspended in 1 mL medium, and counted on a Countess 3 Automated Cell Counter (Invitrogen, AMQAX2000). To generate single cell derived clones, seeding densities were selected to maintain post-irradiation viability and to provide sufficient spacing for discrete colony growth without merging. Cells were irradiated using RadSource RS-2000 X-ray irradiator at pre-defined doses. Cell cultures were then maintained under standard culture conditions with routine medium exchanges until colonies reached the predefined picking size (≈8 mm diameter). Colonies from control and 2 Gy dishes reached picking size within 8 days, whereas 12 Gy colonies required ∼12 days. Colonies were picked with glass cloning cylinders (MilliporeSigma C1059; 8 mm diameter × 8 mm height), trypsinized, transferred to 6-well plates for expansion, and harvested at ∼1 × 10□ cells per clone for DNA extraction.

### Clonogenic Assay

Cells were plated in 15 cm dishes and pre-defined number of cells per dish as above and irradiated at 0, 2, 4, 8, or 12 Gy. When colonies reached 10000 cell size threshold, we stained them with 0.5% (w/v) crystal violet and counted the number of colonies of at least 10000 cells to calculate the surviving fraction under each dosage.

### Whole genome sequencing

Genomic DNA was isolated with the PureLink Genomic DNA Kit (Invitrogen) per the manufacturer’s instructions. DNA concentration was measured by Qubit fluorometric quantification, and samples were stored at −20 °C prior to whole-genome sequencing (WGS). In total, 20 control, 20 (2 Gy), and 20 (12 Gy) clones were submitted for library preparation and Illumina WGS, with a median coverage of ∼30×.

### Sequence Data analysis

Paired-end reads were aligned to the human reference genome (GRCh38, primary assembly) with BWA-MEM v0.7.12. Post-alignment processing followed the Broad Institute Genomics Platform pipeline and GATK Best Practices, with workflow details in ^26,27^, including coordinate sorting, duplicate removal, extraction of discordant reads, ReadCounts generation, haplotype-level allelic coverage estimation, haplotype-specific DNA copy number calculation, and genome-wide coverage visualization. Chromosomal rearrangement junctions were identified from discordantly mapped read pairs using a previously described algorithm^26^.

### De-Novo variant detection

We used GATK HaplotypeCaller (v4.1.2.0) to call short variants in 60 clones. To reduce alignment-related errors and amplification artifacts that do not generate chimeric DNA, we applied stringent post-calling filters. In addition to the default filters, we removed reads with mapping quality (MAPQ) < 10, reads with excessive soft/hard clipping (--filter-too-short 50), and reads with discordant or split alignments. Variant calls supported by fewer than four reads were excluded.

For per-clone quantification of de novo SNVs and indels, variants present in exactly one sample (genotype 0/1 or 1/1) and absent from the remaining samples were classified as de novo, whereas variants observed in over half of the samples were classified as ancestral. To minimize both false positives and false negatives, tetraploid samples were excluded from quantification, under the assumption that whole-genome duplication (WGD) in those clones occurred prior to irradiation. Tetraploid clones were excluded because the expected VAF for heterozygous de novo SNVs in a tetraploid background (≈0.25) overlaps the single-cell artifact peak (≈ 0.20), increasing the risk of misclassifying artifacts as true variants. Moreover, tetraploidy can lead to overestimation the detection sensitivity by increasing the number of ancestral calls due to WGD.

Using copy-number profiles, we identified sets of tetraploid and diploid samples: tetraploids frequently show subclonal arm-level losses across multiple chromosomes, whereas diploids typically exhibit clonal segmental loss (CN 1 to 0). Consistent with ploidy, the de novo VAF distributions peak near 0.25 in tetraploids and near 0.5 in diploids. To avoid confounding from tetraploidy—including flat total CN ≈4—we retained only samples whose de novo VAF distributions showed a clear ∼0.5 peak. After filtering, we retained 9 controls, 5 samples irradiated at 2 Gy, and 14 samples irradiated at 12 Gy for quantification.

We retained heterozygous calls only if the variant allele fraction (VAF) was ≥ 0.32, based on the expectation that true de novo heterozygous variants from diploid samples cluster near VAF ≈ 0.5; VAF cutoff was chosen at the local minimum separating the ∼0.50 (true variant) and ∼0.20 (artifact) peaks (see Supplementary Fig. 2). This filter improves specificity but can increase false negatives. To account for this, we estimated detection sensitivity by using RPE-1 ancestral variants as a reference set and measuring the fraction recovered after applying the VAF ≥0.32 threshold; De novo counts were then calculated as the observed de novo counts divided by sensitivity, yielding the estimated true counts after adjusting for sensitivity.

To minimize chromosome-specific clustering bias, we quantified variant frequency only on disomic (copy-number–neutral) chromosomes. For each disomic chromosome, frequency was calculated as the number of variants divided by chromosome length (Mbp). Short chromosomes (chr15–chr22) and chromosome X were excluded; most clones exhibited X-inactivation.

### Mutation Signature analysis

After extracting true de novo variants, we performed mutational signature analysis on these variants using SigProfilerExtractor (v1.1.23; Python 3.9.19) with default parameters. We then stratified samples into three groups—control, 2 Gy, and 12 Gy—and generated mutation spectra for each group. The mutation signatures extracted were refit by non-negative least square to Catalogue of Somatic Mutations in Cancer (COSMIC) v3 reference^37^ in this tool.

---

**Supplementary Fig. S1.**
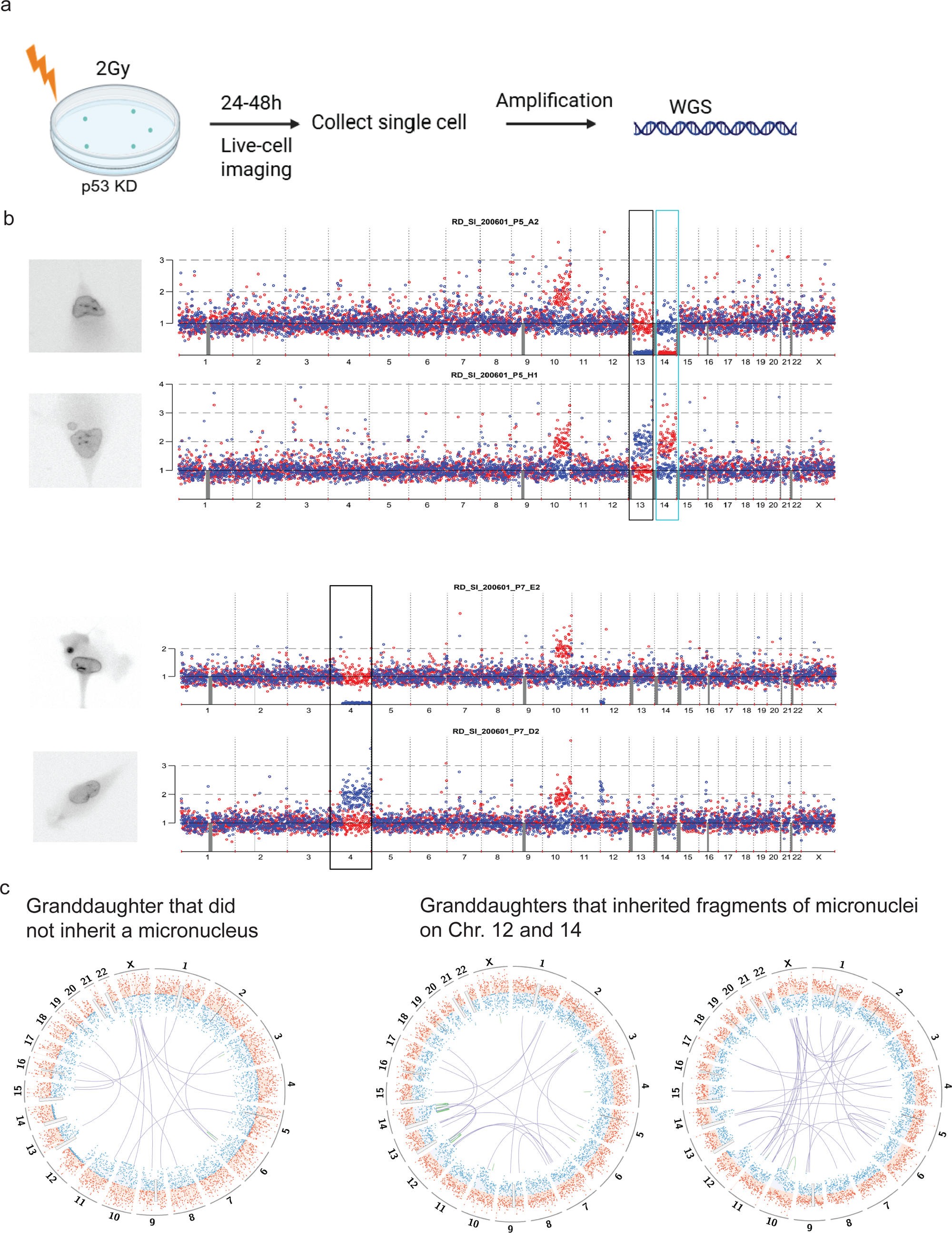
Radiation-induced micronuclei (MN) and chromosome bridges produce genomic consequences comparable to abnormal nuclear structures induced by other perturbations. (**a**) Schematic of the experimental approach. RPE-1 p53-null cells expressing GFP-BAF and PCNA-mCherry were plated 24 hours before irradiation at 2 Gy. Time-lapse live cell imaging started immediately after irradiation and continued for 24–48 h. Snapshots were acquired at 15-minute intervals. Following progression through mitosis, daughter cells that formed MN and/or chromosome bridges were isolated for whole-genome amplification and WGS. (**b**) Representative live-cell imaging snapshots of an isolated daughter cells with MN/bridge and its sister pair (*left*) and corresponding genome-wide copy-number profiles (90-kb bins) (*right*). Blue and red dots denote haplotype-resolved copy number for the two homologs. The boxed region highlights a chromosome exhibiting reciprocal copy-number changes between the two daughter cells (**c**) CIRCOS plots of the haplotype-resolved DNA copy number (90-kb bins) for representative set of granddaughter cells. Three out of 4 granddaughter pairs were successfully isolated. The missing one came from a daughter that did not inherit a micronucleus. Blue and orange dots denote haplotype-specific copy number for the two homologs. Green links indicate intrachromosomal rearrangements and dark blue links indicate interchromosomal rearrangements.

**Supplementary Fig. S2.**
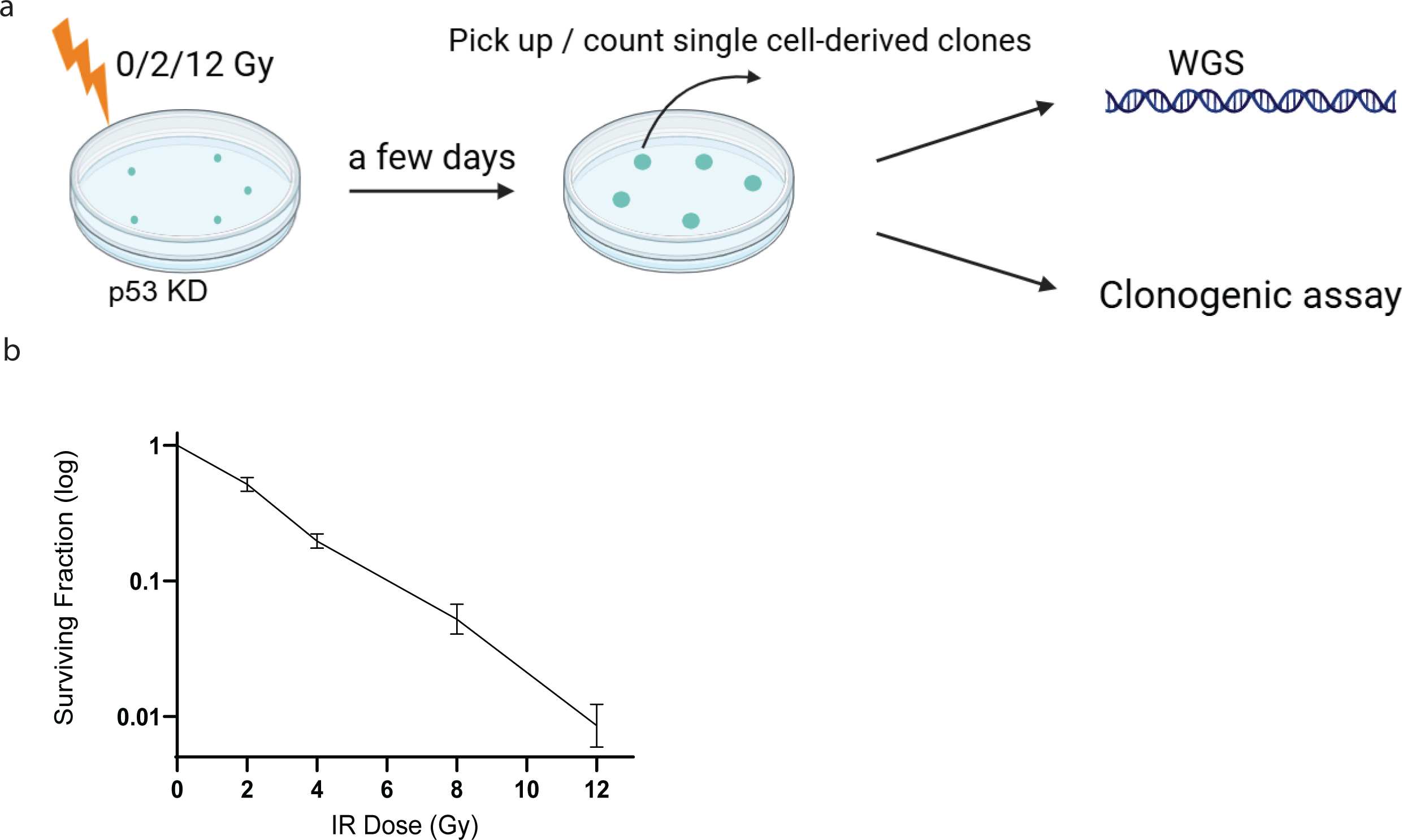
Quantification of clonogenic survival after irradiation. (**a**) Experiment schematic. RPE-1 p53-null cells were plated 1 day before irradiation at 0/2/12 Gy. Half of the plates were used for clonogenic assay, while the other half was used for cell picking for WGS. Once the clones reached 5000 cells on average, they were counted and isolated for WGS. For clonogenic assay, clones that reached at least 10000cells at each dose level were counted. (**b**) Quantification of the percentage of surviving clones after irradiation at 0/2/12 Gy plotted on logarithmic scale. Survival fractions were normalized to the percentage of cells proliferating under 0 Gy. Graphs represent mean ± SEM; n = 3 independent experiments.

